# Structural insights into ubiquitin chain cleavage by *Legionella* ovarian tumor deubiquitinases

**DOI:** 10.1101/2023.01.02.522462

**Authors:** Sangwoo Kang, Gyuhee Kim, Minhyung Choi, Minwoo Jeong, Gerbrand J. van der Heden van Noort, Soung-Hun Roh, Donghyuk Shin

## Abstract

Although ubiquitin is found only in eukaryotes, several pathogenic bacteria and viruses possess proteins that hinder the host ubiquitin system. *Legionella*, a gram-negative intracellular bacterium, possesses an ovarian tumor (OTU) family of deubiquitinases (Lot DUBs). Herein, we describe the molecular characteristics of Lot DUBs. We elucidated the structure of LotA OTU1 domain and revealed that entire Lot DUBs possess a characteristic extended helical lobe (EHL) that is not found in other OTU-DUBs. The structural topology of EHL is the same throughout the Lot family, and it provides an S1′ ubiquitin-binding site. Moreover, the catalytic triads of Lot DUBs resemble those of the A20-type OTU-DUBs. Furthermore, we revealed a unique mechanism by which LotA OTU domains cooperate together to distinguish the length of the chain and preferentially cleaves longer K48-linked polyubiquitin chains. The LotA OTU1 domain itself cleaves K6-linked ubiquitin chains, while it is also essential for assisting the cleavage of longer K48-linked polyubiquitin chains by the OTU2 domain. Thus, this study provides novel insights into the structure and mechanism of action of Lot DUBs.

## INTRODUCTION

Protein ubiquitination, a post-translational modification (PTM), regulates various cellular events in eukaryotes (Yau & Rape, 2016); thus, the disruption of the ubiquitination system leads to severe problems in humans, including cancer, Parkinson’s disease, and neurodegeneration (McNaught *et al*, 2001; Schmidt *et al*, 2021; Swatek & Komander, 2016; Tomaskovic *et al*, 2022). The canonical addition of ubiquitin is tightly regulated by the activating (E1), conjugating (E2), and ligating enzymes (E3) (Dikic & Schulman, 2022; Komander & Rape, 2012; Mukherjee & Dikic, 2022; Popovic *et al*, 2014; Swatek & Komander, 2016; Yau & Rape, 2016). Proteins can be ubiquitinated by monoubiquitin or polyubiquitin chains, and the linkages are formed via the seven lysine residues or the N-terminal methionine residue of ubiquitin (Komander & Rape, 2012; Mattiroli & Sixma, 2014; Swatek & Komander, 2016; Tomaskovic *et al*., 2022). Different linkage types of ubiquitin polymers are involved in different cellular events, such as mitophagy and DNA damage repair (K6- and K27-linked chains), proteasomal degradation (K11- and K48-linked chains), innate immunity (K27-, K33-, and K63-linked chains), cell cycle regulation (K29-linked chains), protein trafficking (K33- and K63-linked chains), and the NF-kB signaling pathway (M1-linked chains) (Swatek & Komander, 2016).

Thus, tight regulation of the proper level of ubiquitination is crucial for cellular viability. Approximately 100 deubiquitinases (DUBs) have been reported in humans; they cleave the isopeptide bond between ubiquitin and the target substrates and counteracts to the E3-ligases (Clague *et al*, 2019; Komander *et al*, 2009). Based on their structure and underlying mechanisms, DUBs are classified into seven different families namely, ubiquitin-specific protease (USP), ovarian-tumor (OTU) domain, Jab1/Mov34/Mpr1Pad1 N-terminal domain (JAMM [MPN]), Machado–Joseph domain (MJD [Josepin]), ubiquitin C-terminal hydrolases (UCH), Zinc Finger with UFM1 specific peptidase domain protein (ZUSFP/ZUP1), and motifs interacting with the novel ubiquitin-containing novel DUB family (MINDY) (Abdul Rehman *et al*, 2016; Clague *et al*., 2019; Haahr *et al*, 2018; Hermanns *et al*, 2018; Hewings *et al*, 2018; Kwasna *et al*, 2018). Except for zinc-containing metalloprotease JAMM (MPN), other DUBs are cysteine proteases (USP, OTU, MJD [Josephin], UCH, ZUSFP, and MINDY). Among these, the OTU family DUBs exhibit ubiquitin-linkage specificity (Mevissen *et al*, 2013; Mevissen & Komander, 2017; Mevissen *et al*, 2016). For example, OTUD1 and OTULIN specifically cleave K63-linked (Mevissen *et al*., 2013) and M1-linked chains (Keusekotten *et al*, 2013), respectively. OTU family DUBs have a structurally similar OTU domain for catalysis, but additional S1 and S2 binding sites, ubiquitin-binding domains (UBDs), and sequence variations at the His loop and variable (V) loop affect their linkage specificity (Mevissen *et al*., 2013).

The ubiquitination system is exclusive to eukaryotes; thus, invading bacteria and viruses have evolved survival strategies to hijack the host ubiquitination system during infection (Lin & Machner, 2017; Wimmer & Schreiner, 2015), as their genomes encode ubiquitin ligases or DUBs that redirect or eliminate host signals. For example, *Salmonella typhimurium* contains the HECT type E3 ligase SopA (Diao *et al*, 2008; Fiskin *et al*, 2017) and *Legionella pneumophila* contains LegU1 and LubX, which are similar to the F-box- and U-box-containing E3 ligases, respectively (Ensminger & Isberg, 2010; Kubori *et al*, 2008; Quaile *et al*, 2015). Not only E3-lgases but also pathogen-encoded DUBs can influence host cellular processes, such as immune response, autophagy, and morphology (Mesquita *et al*, 2012; Pruneda *et al*, 2018; Wan *et al*, 2019).

*Legionella pneumophila*, the causative agent of Legionnaire’s disease, has developed numerous mechanisms of hijacking or inhibiting host ubiquitin signaling. Through defective organelle trafficking/intracellular multiplication (Dot/Icm) type IV secretion system, *L. pneumophila* secretes more than 300 effector proteins into the cytoplasm of the host cell, thereby regulating amongst others host ubiquitin signals (Qiu & Luo, 2017; Segal *et al*, 2005). A representative example is the SidE family ligases (SdeA, SdeB, SdeC, and SidC) of *Legionella* that mediate an unconventional phosphoribosyl (PR) serine ubiquitination system (Bhogaraju *et al*, 2019; Bhogaraju *et al*, 2016; Black *et al*, 2019; Kalayil *et al*, 2018; Qiu *et al*, 2016; Shin *et al*, 2020b). More recently, several studies identified human OTU-like DUBs from Legionella and named them *Legionella* OTU-like DUBs (Lot). Lots include LotA (Lpg2248/ Lem21), LotB (Lpg1621/ Ceg23), LotC (Lpg2529/ Lem27), and LotD (Lpg0227/ Ceg7) (Ashida *et al*, 2014; Kubori *et al*, 2018; Shin *et al*, 2020a). Similar to human OTUs, Lot DUBs exhibit linkage specificity. LotB and LotC prefer the K63- and K48-linkage, respectively. In contrast, Ceg7 cleaves K6-, K11-, K33-, K48-, and K63-linkage (Kitao *et al*, 2020; Schubert *et al*, 2020; Shin *et al*., 2020a). Interestingly, LotA comprises two OTU-like domains with two catalytic cysteine residues (C13 and C303). The first OTU domain of LotA (LotA OTU1) exclusively cleaves the K6-linkage, whereas the second OTU domain (LotA OTU2) cleaves K48- and K63-linkages (Kubori *et al*., 2018; Takekawa *et al*, 2022).

Despite various efforts, however, the conserved molecular mechanism of this unique pathogenic DUBs has not been clearly understood. In this study, therefore, we investigated the structure and molecular mechanism of Lots. We found that Lots share a unique structural topology that is not found in any other OTUs. In addition, the catalytic triads of Lots were revealed to have differences compared to other OTUs. Moreover, we also provide molecular insights into how Lots distinguishes specific Ub-linkage types and length of ubiquitin chains.

## RESULTS

### LotA domains exhibit cooperative catalysis

Legionella contains four OTU-deubiquitinases (LotA (Lpg2248/ Lem21), LotB (Lpg1621/ Ceg23), LotC (Lpg2529/ Lem27), LotD (Lpg0227/ Ceg7)) that exhibit distinct ubiquitin chain specificity. Among them LotA comprises two OTU domains (LotA OTU1 and LotA OTU2, Fig 1A). Previous reports defined the linkage specificity of each OTU domain of LotA(Kubori *et al*., 2018; Takekawa *et al*., 2022). LotA OTU1 preferentially cleaves K6-linked chains, while LotA OTU2 cleaves K48 and K63 chains. Consistent with the previous reports, LotA OTU1 domain (LotA_7–290_) effectively cleaved K6-linked diubiquitin chains (Fig 1B). However, neither LotA OTU2 (LotA_294–544_) nor LotA OTU1_OTU2 (LotA_7–544_) domains cleaved K48- or K63-linked diubiquitin chains to form a covalently linked complex (Fig 1C and D). We further analyzed the K48- or K63-linkage specificity of LotA OTU2 with diubiquitin activity-based probes (Fig EV1A–D). Propargyl (Prg) probes, which have a Prg group at the C-terminus of the diubiquitin chain, can detect S1 and S2 sites of DUBs (Ekkebus *et al*, 2013; Flierman *et al*, 2016; Sommer *et al*, 2013). Vinyl-methyl ester (VME) probes, which have a VME reactive group between the two ubiquitin moieties of a diubiquitin chain, were developed to identify S1 and S1′ sites on DUBs (Borodovsky *et al*, 2002; Mulder *et al*, 2014). Similar to the results obtained for the cleavage analysis of diubiquitin chains, none of the LotA OTU domain constructs (OTU1, OTU2, and OTU1_OTU2) reacted with the diubiquitin activity-based probes of K48- or K63-linkage chains (Fig EV1A–D), indicating that the catalytic cysteines of OTU1 (C13) and OTU2 (C303) domains did not react with the chemical warheads.

**Figure 1.**
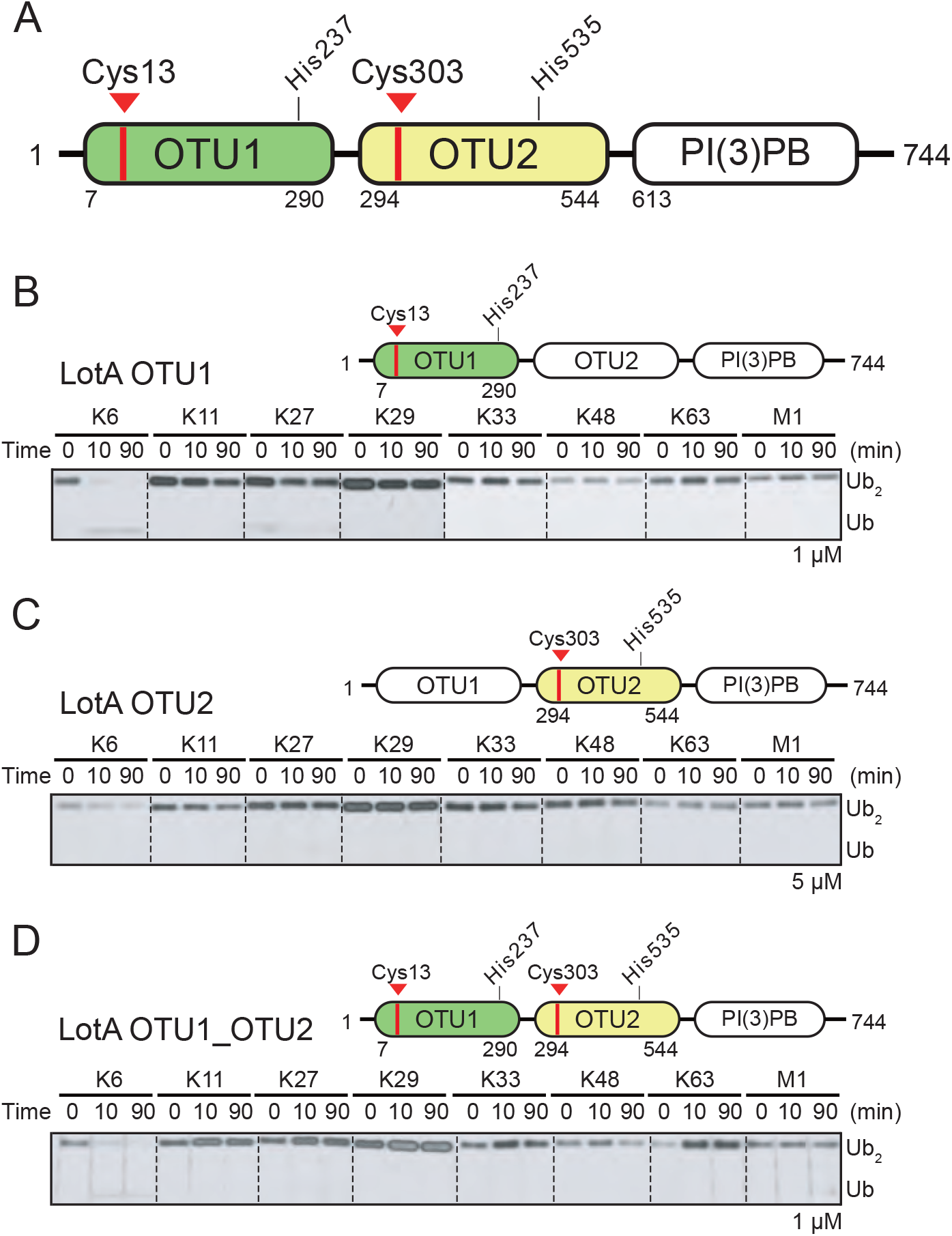
Deubiquitinase activities of both OTU domains of LotA. A. The schematics of *L. pneuomophila* LotA domain architecture. LotA comprises two OTU domains (LotA OTU1, LotA OTU2). Catalytic cysteines of each OTU domain are labeled. B. Di-ubiquitin cleavage assay by LotA OTU1 (LotA_7-290_) against eight different di-ubiquitin linkages. Reactions were quenched at the indicated time-points and analyzed on SDS-PAGE with silver staining. C. Di-ubiquitin cleavage assay by LotA OTU2 (LotA_294-544_) against eight different di-ubiquitin linkages. Reactions were quenched at the indicated time-points and analyzed on SDS-PAGE with silver staining. D. Di-ubiquitin cleavage assay by LotA OTU1_OTU2 (LotA_7-544_) against eight different di-ubiquitin linkages. Reactions were quenched at the indicated time-points and analyzed on SDS-PAGE with silver staining.

Interestingly, all previous studies used longer polyubiquitin chains with more than two ubiquitin residues to elucidate the cleavage of K48- and K63-linked ubiquitin chains by the OTU2 domain (Kubori *et al*., 2018; Takekawa *et al*., 2022). Therefore, we examined the catalytic activity of the LotA OTU2 domain in cleaving K48- or K63-linked tetraubiquitin chains. We determined that the LotA OTU2 domain cleaved both K48- and K63-linked tetraubiquitin chains, whereas the LotA OTU1 domain did not cleave either of the two chains (Fig 2A–C). These results revealed the length-dependent catalytic activity of LotA OTU2, which only cleaves polyubiquitin chains with more than two ubiquitin moieties. In contrast, the K6-linkage specificity of LotA OTU1 was not restricted by the length of polyubiquitin chains, as it cleaved K6-linked diubiquitin chain (Fig 1B).

**Figure 2.**
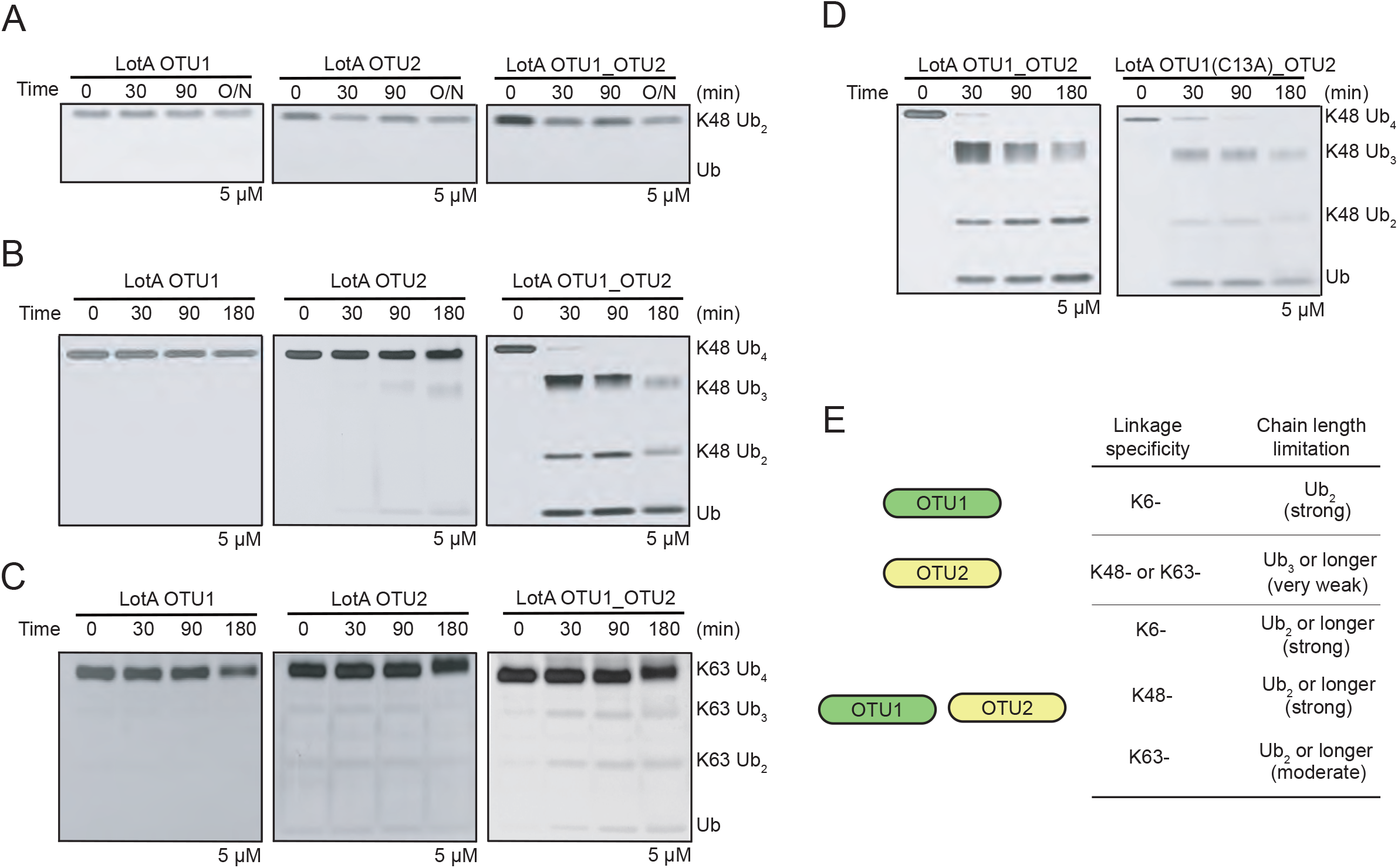
Cooperative catalytic activity of LotA OTU1 and OTU2 domain. A. K48-linked di-ubiquitin cleavage assay by LotA OTU constructs (OTU1, OTU2, OTU1_OTU2). B. K48-linked tetra-ubiquitin chains cleavage assay by LotA OTU constructs (OTU1, OTU2, OTU1_OTU2). Catalytic activity against K48-linked tetra-ubiquitin chains were remarkably enhanced when intact OTU1 and OTU2 domain (LotA_7-544_) was used. C. K63-linked tetra-ubiquitin chains cleavage assay by LotA OTU constructs (OTU1, OTU2, OTU1_OTU2). Catalytic activity against K63-linked tetra-ubiquitin chains was enhanced when intact OTU1 and OTU2 domain (LotA_7-544_) were used. D. K48-linked tetra-ubiquitin chain cleavage assay by wild-type LotA OTU1_OTU2 and LotA OTU1*_OTU2 (C13A) mutant. E. Ubiquitin linkage and length preference of LotA OTU constructs (OTU1, OTU2, OTU1_OTU2).

Surprisingly, the cleavage of K48- and K63-linked tetraubiquitin chains by LotA was remarkably enhanced when intact OTU1 and OTU2 domains (LotA_7-544_) were used for the cleavage assay (Fig 2B–C). While the apparent processing speed of K48-chains is significantly higher than K63-chains, both K48- and K63-chains were cleaved much more with OTU1_OTU2 construct than OTU2 construct, indicating the supportive role of the OTU1 domain in K48- or K63-chain processing. Subsequently, we investigated whether the catalytic activity of the OTU1 domain was required for the cleavage of K48- and K63-linked chains by the OTU2 domain. Thus, we mutated the catalytic cysteine (Cys13) in OTU1 to alanine (LotA OTU1*-OTU2; LotA_7-544_ C13A) and observed that the LotA OTU1*-OTU2 construct retained its catalytic activity and cleaved K48-linked tetraubiquitin chain at the same rate as that of the wild-type (WT; Fig 2D), indicating that the catalytic activity of the OTU1 domain is not necessary for the cleavage of the K48- or K63-polyubiquitin chains, but its presence is crucial for the catalytic activity of the OTU2 domain (Fig 2D and E).

Next, to understand the molecular mechanism underlying the interplay between the OTU1 and OTU2 domains, we analyzed the overall architecture of LotA OTU1_OTU2 (LotA_7-544_) using electron microscopy. Even though the protein size of LotA_7-544_ is on the border of current Cryo-EM technique, we expected the observation of the relative orientation and overall shape. 2D class averages of the negative stained- and cryo-images displayed heterogeneous classes with two distinct globular domains (Fig 3A and EV2). We then reconstructed cryo-EM map at ∼11 Å (cryo-EM, Fig 3B and E and Fig EV2), showing two converged densities of the LotA_7–544_. Although the resolution was not sufficient to model the precise atomic details of both OTU domains, the overall dimensions suggested that the two OTU domains were not separated but aligned in proximity (Fig 3B; Supplementary table 1). While our cryo-EM work was underway, a crystal structure of LotA OTU1_OTU2 was deposited (PDB: 7W54). Intriguingly the crystal structure contains completely different conformations of LotA molecules in the asymmetric unit (ASU). Superimposition of the OTU1 domain of both conformers in the ASU exhibited approximately 180° rotation of the OTU2 domain, indicating that the relative orientation of the OTU1 and OTU2 domains varied in the solution (Fig 3C). We could not fit both high-resolution conformers into our cryo-EM electron density map. Therefore, we hypothesized that the cryo-EM electron density map presented different orientation of OTU1-OTU2 domains with respect to each other and perhaps the domain orientation is variable in solution. These results prompted us to assume that a specific structural arrangement of the OTU1 and OTU2 domains might be required for the recognition and cleavage of the K48- and K63-linked tetraubiquitin chains (Fig 2B–D). To prove that the OTU1 domain structurally assists the OTU2 domain, we deleted a linker region (276–283) between the two OTU domains and restricted the conformational spaces of the OTU domains. We observed that the linker-deletion mutants exhibited slower cleavage of K48-linked tetraubiquitin chains than the WT or C13A mutants (Fig 3D), indicating that structural flexibility between the two domains is required for K48-linked tetraubiquitin recognition and cleavage. Together, our results reveal that the OTU1 domain of LotA specifically cleaves K6-linked polyubiquitin chains without any length limitation, whereas the OTU2 domain processes K48- or K63-linked polyubiquitin chains in the presence of more than two ubiquitin moieties. Moreover, our data revealed that the OTU1 domain is not only required for K6-linked polyubiquitin chain cleavage but also assists the OTU2 domain in recognizing and cleaving K48- or K63-linked polyubiquitin chains.

**Figure 3.**
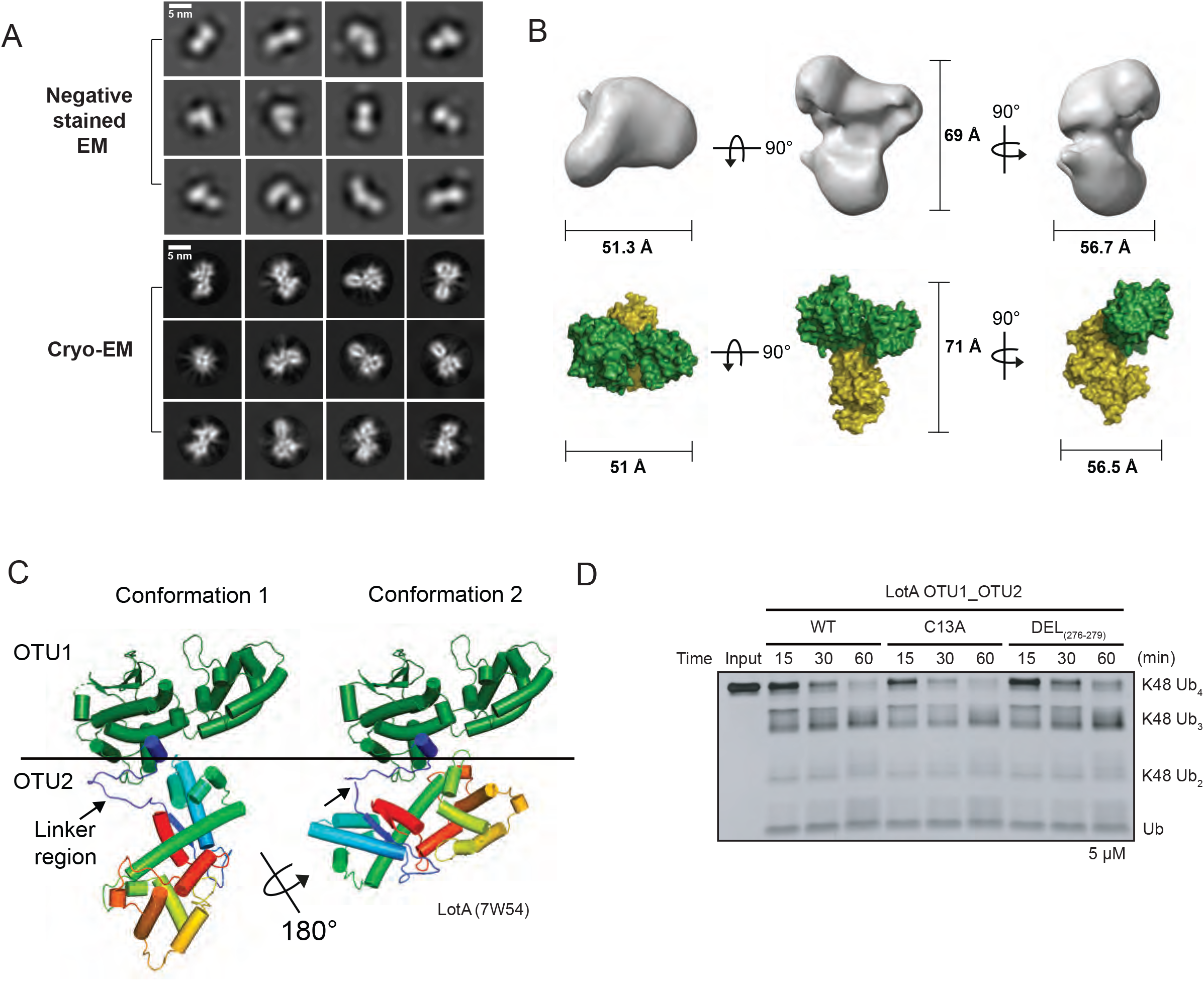
Structure of LotA OTU1_OTU2. A. Selected 2D classes of negative stained (upper) and Cryo-EM (down) LotA OTU1_OTU2. Scale bar represented 5 nm. B. LotA 7-544 cryo-EM model orientated X, Y, Z axis. Each distance is described as angstrom (Å)(upper). Proposed OTU1 and OTU2 orientation of LotA 7-544 (down) using crystal structure (PDB: 7w54). C. Crystal structure of LotA OTU1_OTU2 (7W54). Two completely different conformations of LotA molecules are observed in the asymmetric unit (ASU). D. DUB assay comparing cleavage of K48-linked tetra-ubiquitin chains by wild-type LotA OTU1_OTU2, LotA OTU1*_OTU2 (C13A) mutant, and linker deletion (residues 276-279) mutant of LotA OTU1_OTU2.

### Crystal structure of the OTU1 domain of LotA

To elucidate the mechanism underlying the cleavage of K6-linked polyubiquitin chains by the LotA OTU1 domain, we successfully obtained the crystal structure of the LotA OTU1 domain. Based on secondary structure analysis, we designed a LotA OTU1 construct spanning residues 7–290 (LotA_7–290_). The LotA_7–290_ crystal diffracted at 1.54 Å resolution, and we determined the structure by molecular replacement using the structure predicted by the AlphaFold Protein Structure Database as a template model (Fig 4A and EV5A) (DiMaio *et al*, 2011; Evans & McCoy, 2008; Jumper *et al*, 2021; Scapin, 2013). The crystal belonged to the tetragonal space group P4_1_2_1_2, with one molecule per ASU. The final refined model of LotA_7-290_ contained 270 residues of LotA (Ala7 to Thr276) and three amino acids from the glutathione-S-transferase (GST) expression tag (Table 1).

**Figure 4.**
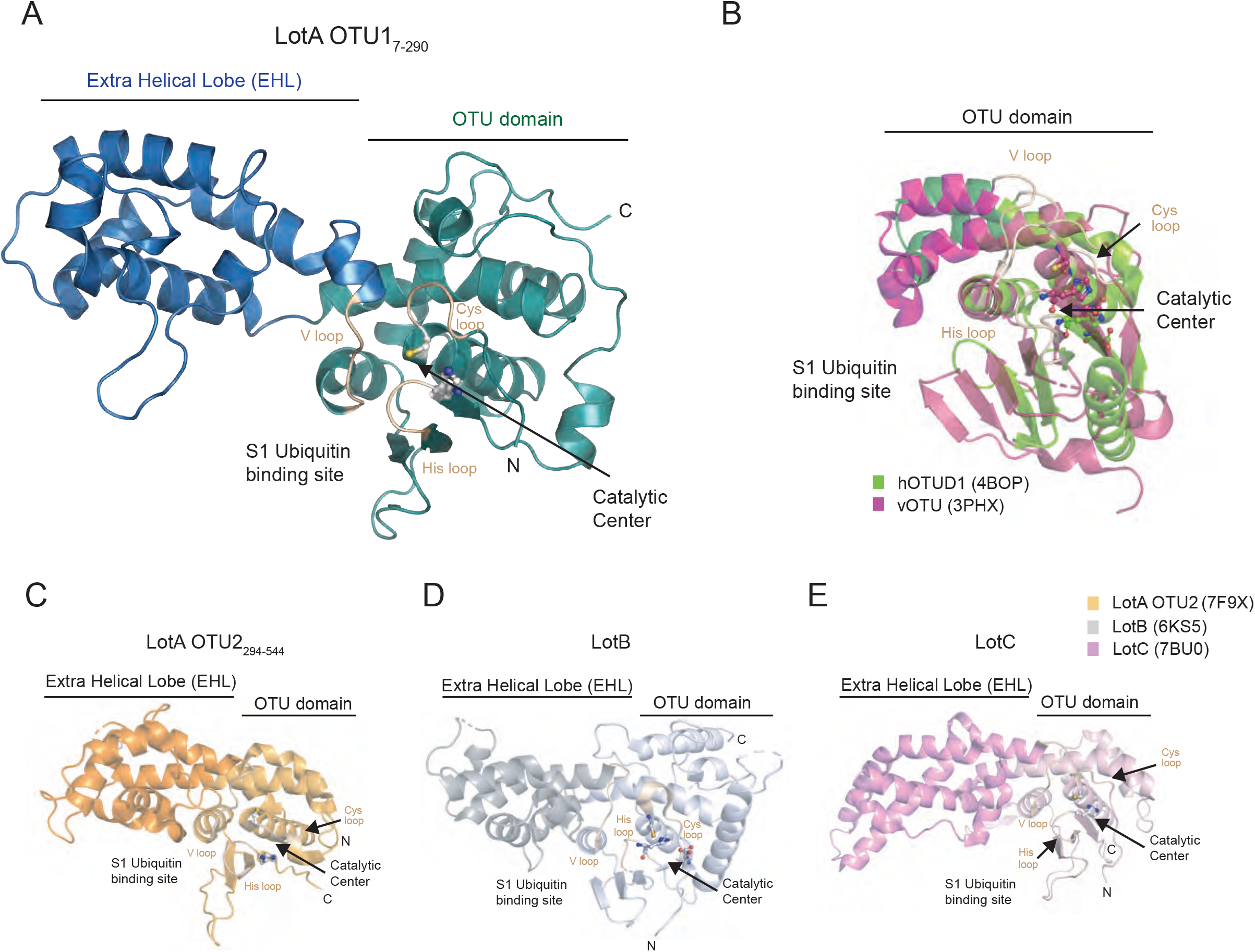
Structural comparison of Legionella OTU deubiquitinases. A. X-ray Crystal structure of LotA OTU1 (LotA_7-290_) diffracted to 1.54 Å determined by molecular replacement (Deposited PDB ID: 8GOK). Catalytic cysteine and histidine of LotA OTU1 are shown as ball and stick model and Cys loop, His loop, and V loop are highlighted in wheat color. EHL domain and catalytic OTU domain are labeled. B. Structural comparison of hOTUD1 (4BOP, green) and vOTU (3PHX, pink). Structures were aligned by their core OTU domain. C. Crystal structure of LotA OTU2 (7F9X). Catalytic cysteine and histidine of LotA OTU2 are shown as ball and stick model. EHL domain and catalytic OTU domain are labeled. D. Crystal structure of LotB (6KS5). Catalytic triad residues of LotB are shown as ball and stick model. EHL domain and catalytic OTU domain are labeled. E. Crystal structure of LotC (7BU0). Catalytic triad residues of LotC are shown as ball and stick model. EHL domain and catalytic OTU domain are labeled.

**Table 1.**
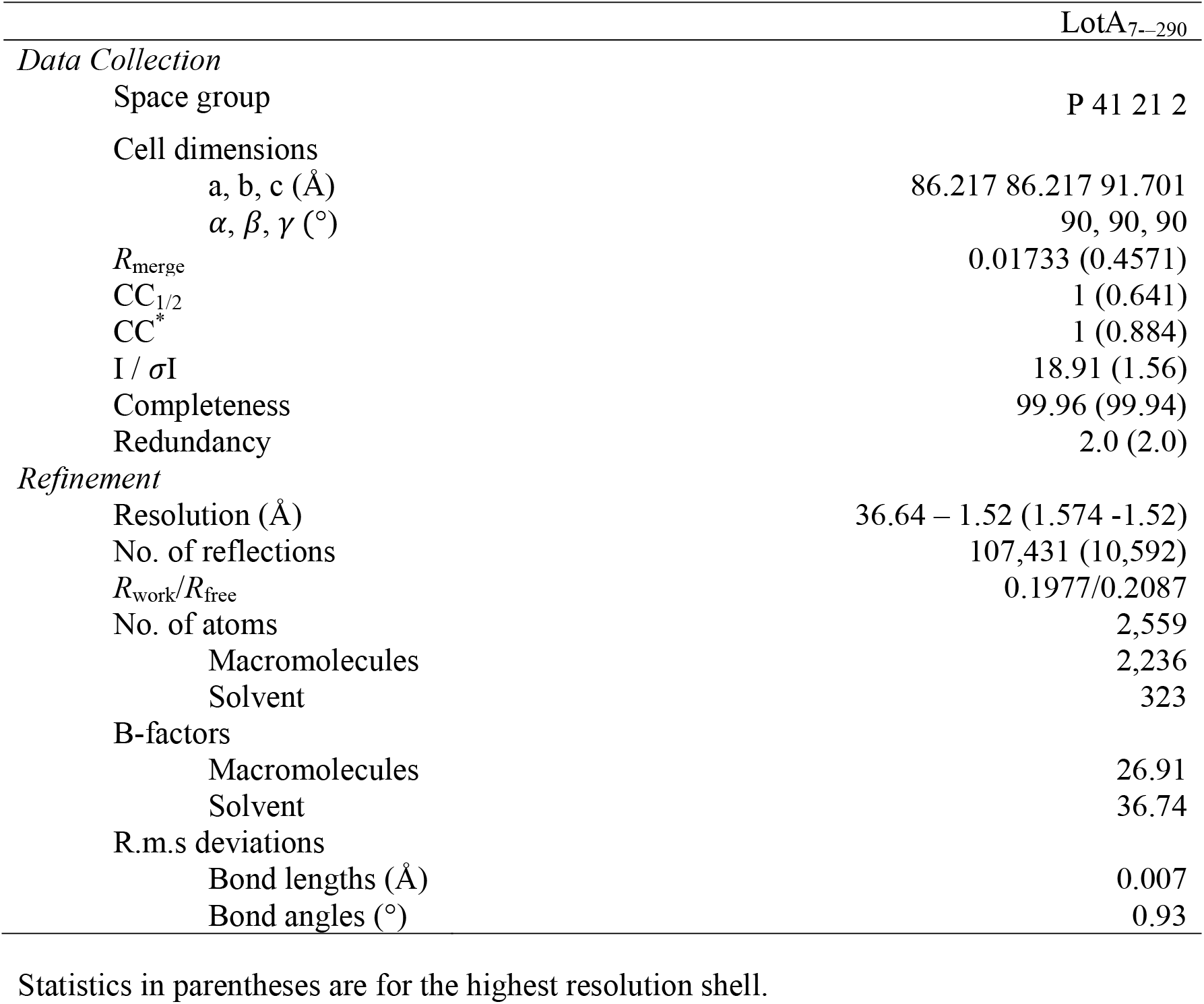
X-ray data collection and refinement statistics.

The overall structure of LotA_7–290_ shows a dumbbell-like fold harboring a typical OTU fold (residues 7–90, 200–276) and an EHL part (residues 91–199) (Fig 4A and B, and EV3A) (Mevissen *et al*., 2013; Shin *et al*., 2020a). The OTU domain of LotA_7-290_ comprises eight α-helices (α1–α5 and α13–α15) and a β-sheet with three β-strands (β1–β3). The catalytic center of the OTU domain is composed of a catalytic cysteine residue (Cys13) containing α1 (residues 13–27) and a β-sheet containing a histidine residue (H237). The EHL of the LotA OTU1 domain is composed of only α-helices and is located before the conserved V-loop and near the S1 ubiquitin-binding site, suggesting a putative role of the EHL in K6-linked polyubiquitin selectivity of LotA OTU1 (Fig 4A, C-E) (Kubori *et al*., 2018; Shin *et al*., 2020a). To examine whether the extra insertion of EHL domain of LotA_7–290_ resembled any known structures, we performed a structural similarity analysis (SSA). The Dali-SSA server found that the structure of LotA_7-290_ resembles that of previously known Lot structures (LotA_OTU2 (PDB: 7f9x), LotB (PDB: 6ks5), and LotC (PDB: 7bu0)), indicating that the insertion of the EHL domain is a unique structural feature of the Lot family and distinguishes it from other DUB families.

To further characterize the EHL domain of the Lot DUB family, we compared the topology of the EHL domains of all known Lot DUBs (Kubori *et al*., 2018; Liu *et al*, 2020; Ma *et al*, 2020; Shin *et al*., 2020a; Takekawa *et al*., 2022). Our analysis revealed that the EHL domains of all Lot family members have similar topologies (Fig 5A-B). The EHL domain contains a roof-like fold, which is composed of three helices before the V loop, and an amphipathic alpha helix is present below the roof folding. The hydrophobic region of the amphipathic helix is in contact with the roof helices, whereas the hydrophilic surface is exposed to the surface. To validate whether this structural feature of EHL was also present in LotD (Ceg7), the AlphaFold-predicted structure of LotD was analyzed (Fig EV3C). Interestingly, the EHL of LotD has the same topology as that of the other Lots. These results suggest that the topology of the EHL is unique to the entire Lot DUB family and can be considered a standard characteristic of Lots.

**Figure 5.**
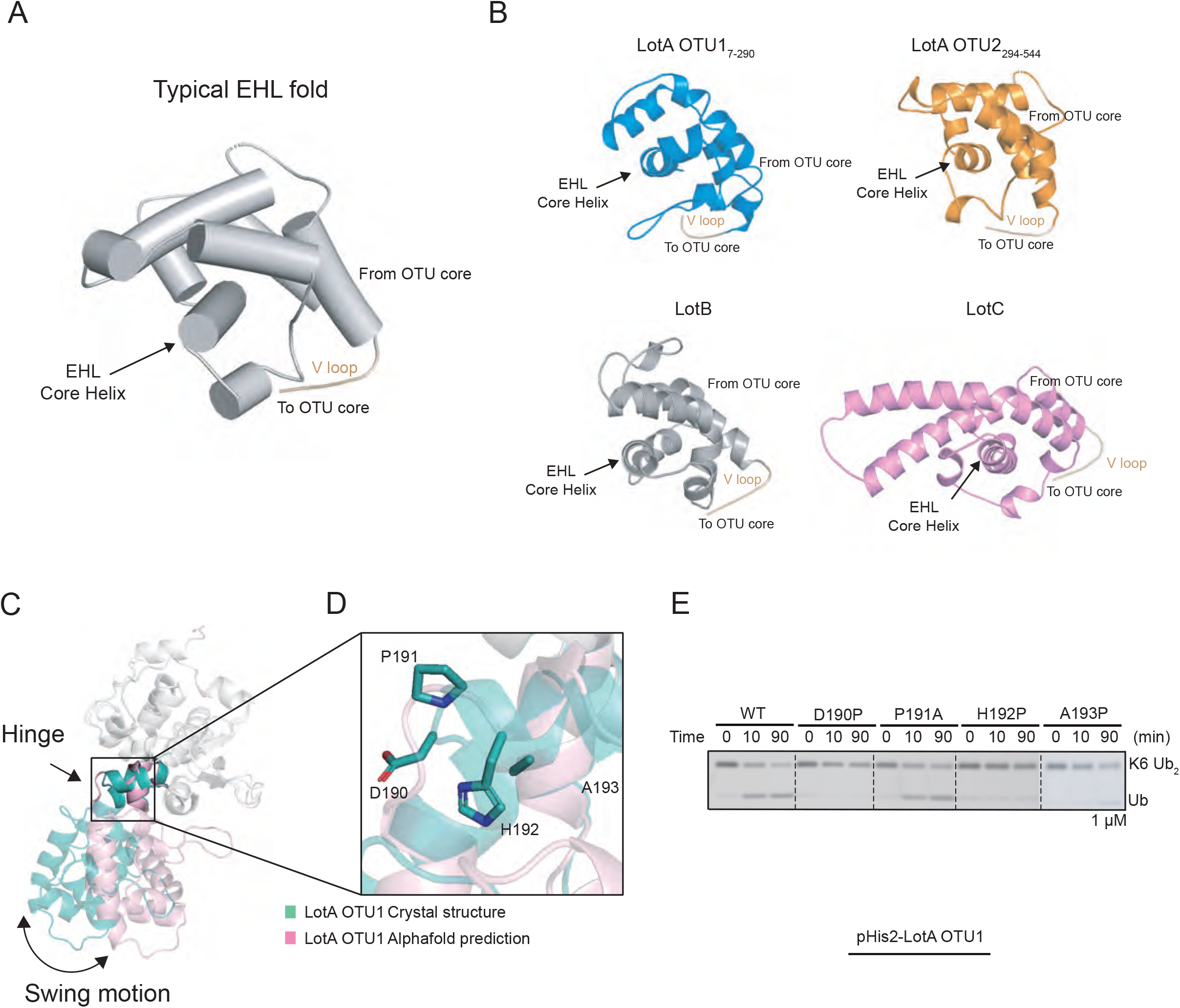
Structural analysis of the EHL domain of Lots. A. Typical EHL fold architecture of LOT DUB family. B. Structure of EHL domain of LOT DUB family. The EHL domain of LOT DUB family shared roof-like folding. C. Structural comparison of LotA OTU1 crystal structure (cyan) and LotA OTU1 Alphafold prediction (pink). OTU domain of two structures are superimposed and relative orientation of EHL domain to OTU domain is presented. D. Close-up view of hinge region of LotA OTU1. Four amino acids (residues 190-193) in the hinge region between EHL and OTU domains are shown as stick model. E. DUB assay against K6-linked di-ubiquitin with wild-type and the hinge mutants of LotA OTU.

### Conformational change in EHL is required for K6-linked polyubiquitin chain cleavage by LotA

During the structure determination of the LotA_7–290_ with molecular replacement using the AlphaFold-predicted structure as a template model, we could not obtain a successful solution. However, when the OTU and EHL domains were manually divided and used as two individual templates, we obtained the phase information and successfully determined the structure of LotA OTU1_7–290_ (Fig EV5A). This led us to explore the conformational diversity of the EHL of the OTU1 domain. Moreover, previous studies on LotC have revealed that binding of ubiquitin to OTU induces conformational changes in the EHL domain (Fig EV5B) (Liu *et al*., 2020; Shin *et al*., 2020a). To examine whether this ubiquitin-binding-dependent conformational change was also required for the catalytic activity of LotA OTU1, we designed an experiment to restrict the conformational flexibility of the EHL domain. Based on structural comparison, four amino acids (N190 to A193) in the hinge region between the EHL and the OTU domains were chosen for examination. We assumed that substitution of each of these amino acids with proline restricted the relative movement of EHL to the OTU domain. Indeed, the cleavage of K6-linked diubiquitin chain was completely abolished in all three proline mutants (D190P, H192P, and A193P) (Fig 5C-E). We also introduced an alanine mutation at P191 in the hinge region to examine whether it induces the EHL domain to a specific orientation with respect to the OTU1 catalytic domain. However, the catalytic activity of the P191A mutant was similar to that of the wild-type OTU1, suggesting that P191 was not responsible for placing EHL in a specific orientation with respect to the OTU domain. Thus, our results suggest that the conformational flexibility of the EHL domain of OTU1 is crucial for K6-linked diubiquitin chain cleavage, and this conformational change in the EHL domain upon recognition of the ubiquitin moiety could be a unique characteristic of the Lot DUB family.

### Catalytic triad of Lot DUBs resembles that of A20 OTU family

The catalytic sites of cysteine proteases, including OTU-DUBs, harbor three residues (catalytic triads, cysteine, histidine and an acidic amino acid;Asp or Glu) (Baker *et al*, 1993; Buller & Townsend, 2013; Mevissen *et al*., 2013; Verma *et al*, 2016). Both cysteine and histidine residues actively participate in cleaving the peptide bonds. The catalytic cysteine residue attacks the carbonyl carbon of the scissile peptide bond, and this step requires a histidine residue that accepts a hydrogen from the thiol group of the cysteine residue. The third residue of the catalytic triad is acidic (aspartic acid or glutamic acid), and its primary role is to stabilize the charge on the imidazole ring of the histidine residue during catalysis. Although both cysteine and histidine residues are indispensable for peptide bond cleavage and are highly conserved in the cysteine protease family, the acidic residue is not required for the activity of some cysteine proteases, which have catalytic dyads (Elsasser *et al*, 2017; Ferreira *et al*, 2021; Nakagawa, 2013; Ramos-Guzman *et al*, 2020). Recent studies on DUBs from different bacteria have suggested variations in the catalytic triad (Akutsu *et al*, 2011; Nakagawa, 2013; Pruneda *et al*, 2016; Schubert *et al*., 2020; Shin *et al*., 2020b; Takekawa *et al*., 2022). However, the catalytic residues of Lot DUBs have not been clearly defined. For instance, although both LotB and LotC possess an acidic amino acid residue (D21 and D17, respectively) present N-terminally from the catalytic cysteine residue (C29 and C24, respectively), LotA_OTU2 and LotD have an asparagine or a threonine residue present C-terminally from the catalytic cysteine residues.

To elucidate the catalytic residues of Lot DUBs, we analyzed the structures of Lot DUBs. Because misaligned catalytic triads are often observed in the crystal structures, we also compared the structures predicted using the AlphaFold database (Fig 6A). Previous studies have reported catalytic triads of both LotB and LotC, where the arrangement of triads resembles the catalytic site of A20 or Cezanne-type OTUs (Komander & Barford, 2008; Ma *et al*., 2020; Mevissen *et al*., 2016; Shin *et al*., 2020a). Both LotB and LotC have an acidic residue located N-terminally from the catalytic cysteine residue, which differs from typical OTU-DUBs, which have an acidic residue located two amino acids C-terminal of the histidine residue on the same beta-strand (His-XX-Acidic residue, Fig 6B). Interestingly, similar to the structures of LotB and LotC, our structural analysis revealed the presence of negatively charged residues toward the N-terminal region from the catalytic cysteine residue in case of other Lot DUBs, including LotA_OTU2 and LotD. Both LotA_OTU2 and LotD exhibited a well-positioned catalytic triad with the histidine base between the catalytic cysteine residue and acidic amino acids (Fig 6A). Consistent with the structural analysis, mutation at D296 (D296A) of LotA OTU2 and D6 (D6A) of LotD reduced the enzymatic activity of LotA_OTU2 and LotD, respectively (Fig 6C). Previous reports have suggested that both LotA OTU2 have a threonine residue at the C-terminal position to the base histidine residue, which is similar to canonical OTU-DUBs (Clague *et al*., 2019; Edelmann *et al*, 2009; Mevissen *et al*., 2013; Schubert *et al*., 2020; Takekawa *et al*., 2022). However, the mutation at T537 position of LotA OTU2 (T537A) did not affect the catalytic activity, suggesting that D296 is N-terminally from the catalytic C303 residue of LotA OTU2 as the acidic residue complementing the catalytic triad of LotA OTU2. These results indicate that the catalytic triads of Lot DUBs belong to the A20-type, which has an acidic amino acid present located N-terminally from the catalytic cysteine residue. We also validated other predicted residues of LotA_OTU2 and LotD, which also comprised catalytic triads, by introducing mutations in the predicted residues (Fig 6C and EV4A). All mutations, indeed, completely abolished the catalytic activity of Lot DUBs. For LotA OTU1, we could not identify the acidic residue present N-terminally from the catalytic cysteine residue. Instead, LotA OTU1 had an asparagine residue (N239) at the C-terminus of the base histidine residue (H237). Mutation N239 (N239A) exhibited less catalytic activity than the WT, and more K6-linked diubiquitin chains were present (Fig 6C), indicating that the catalytic core of LotA OTU1 follows that of typical OTU-DUBs. Further structural analysis of LotA OTU1 revealed two glutamate residues (E235 and E236) located at the N-terminal from the H237 residue and an N239 residue at the two amino acids C-termini of H237. However, mutations in these residues did not affect the catalytic activity. Instead, D10, which is located toward the N-terminus from the catalytic cysteine residue, completely abolished the cleavage of K6-linked diubiquitin chains. Because both the crystal and AlphaFold predicted structures did not exhibit the possibility of the presence of D10 to stabilize H237, and D10 is a part of the oxyanion hole, we assumed that the D10A mutant altered the conformation of the oxyanion hole loop. To test this hypothesis, we introduced a proline mutation close to (T8P) or further away (A7P) from Asp10. Interestingly, the single proline mutation at T8 position also completely diminished the catalytic activity of the LotA, supporting the role of D10 in the formation of an oxyanion hole. Together, these results revealed that Lot DUBs resemble the catalytic triad of A20 OTU, although LotA OTU1 possesses a slightly altered catalytic core compared to other OTU-DUBs.

**Figure 6.**
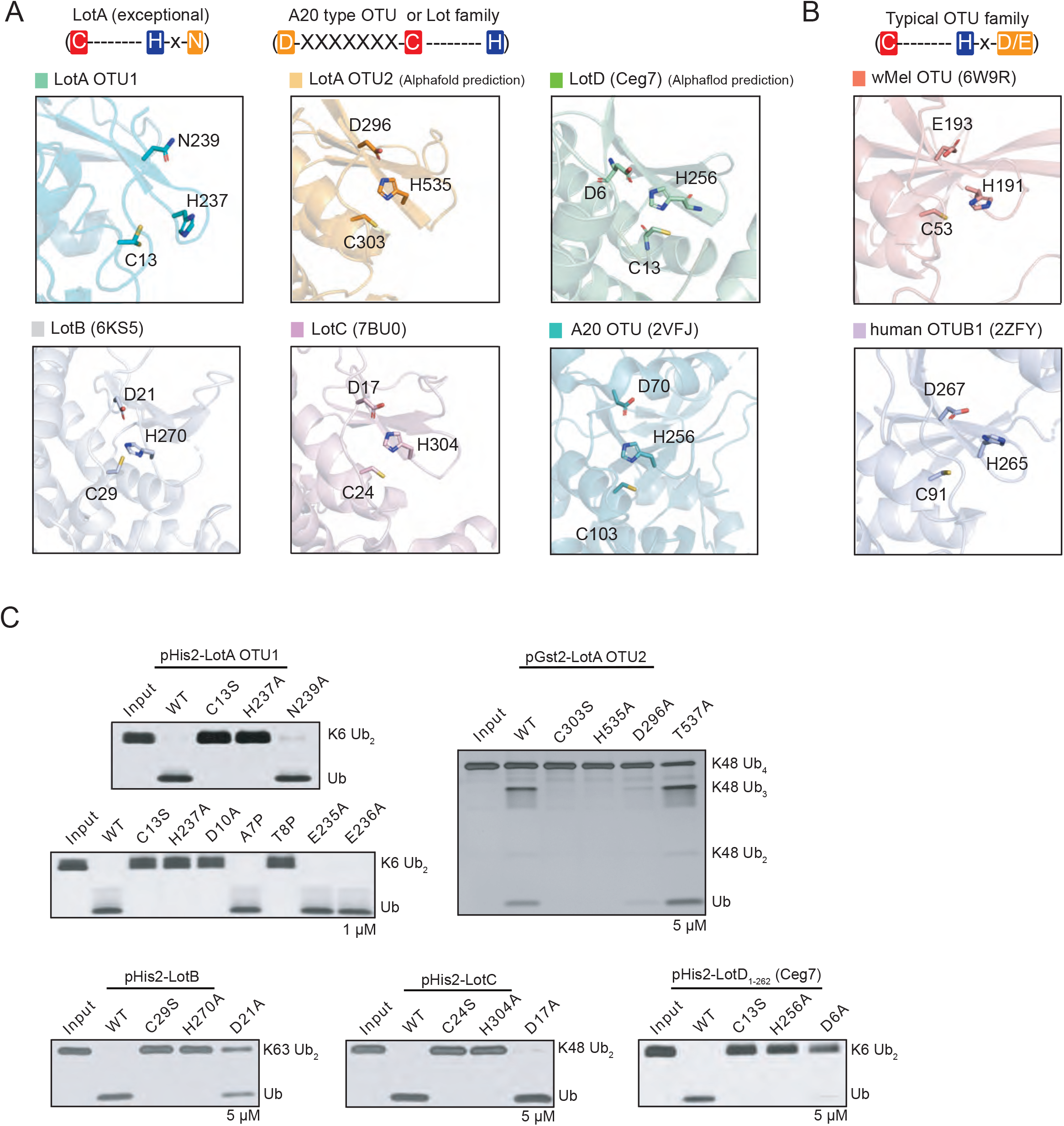
Catalytic triad of LOT DUB family. A. Close-up views of the LotA OTU1, LotA OTU2, LotB (6KS5), LotC (7BU0), LotD, A20 OTU (2VFJ) catalytic site. Catalytic triads are highlighted as stick model. Schematic representation of the order of catalytic triads are also presented. B. Close-up views of the wMel OTU (6W9R), hOTUB1 (2ZFY) catalytic site. Catalytic triads are highlighted as stick model. Schematic representation of the order of catalytic triads are also presented. C. Di-ubiquitin or poly-ubiquitin cleavage assay by LOT DUB family to define catalytic triad residues. The catalytic activity of LOT DUB family wild-type and their mutants were tested. Reactions were quenched at the indicated time-points and resolved by SDS-PAGE with silver staining, respectively.

### Structural elements of S1 ubiquitin-binding site of Lot family

Previous studies on bacterial DUBs, including CE-clan ubiquitin or ubiquitin-like proteases and OTU-DUBs, have reported the key regions that form the S1 ubiquitin-binding site (Akutsu *et al*., 2011; James *et al*, 2011; Mevissen *et al*., 2013; Pruneda *et al*., 2016; Schubert *et al*., 2020). OTU-DUBs include a structurally conserved helix (constant region, CR) and three variable regions (VRs; VR1–3) (Schubert *et al*., 2020). In general, VR1 is a helical arm located underneath the CR-helix, VR2 refers to the edge of the beta-sheets, and VR3 is the beta-hairpin region. Since all Lot family members have an EHL domain, which is located close to the S1-ubiquitin binding site, we investigated the effects of the EHL domain on the S1 ubiquitin-binding site. Structural comparison of Lot DUBs revealed that the position of the Lot EHL domain overlapped with the VR1 of other OTU-DUBs (Fig EV6A). Moreover, previous studies on LotB and LotC have experimentally verified several residues that are crucial for ubiquitin binding (Fig EV6C–E), and these residues are concentrated in the C-terminal helical loop region of the EHL core helix. Interestingly, this helical loop region is also conserved in the crystal structures of LotA OTU1 and OTU2 domains and the predicted structure of LotD, suggesting their putative role in ubiquitin binding (Fig EV3C and EV6B). Based on these observations, we hypothesized that this helical loop region in the EHL domain forms the VR1 in the Lot family (Fig 7A). To test this, we mutated a number of residues in this loop of LotA OTU1 and examined the catalytic activity of these mutants. Surprisingly, eight of the twelve single amino acid mutants were considerably less active in cleaving K6-linked diubiquitin chains compared to the WT (Fig 7B and C), implicating the role of this helical loop as VR1 to form the S1 ubiquitin-binding site in the Lot family. To further characterize the S1 ubiquitin-binding site, we examined other VRs (VR2 and VR3) of LotA OTU1. Two mutations (L214A and E216R) in VR2 exhibited less catalytic activity compared to that of the WT, and mutations in VR3 did not affect the activity, suggesting a less important role of VR3 in the recognition of K6-linked diubiquitin chains. Therefore, our results indicate that the helical loop present toward the C-terminal from the CR is the VR1 of Lot family, and it forms a part of the S1 ubiquitin-binding site.

**Figure 7.**
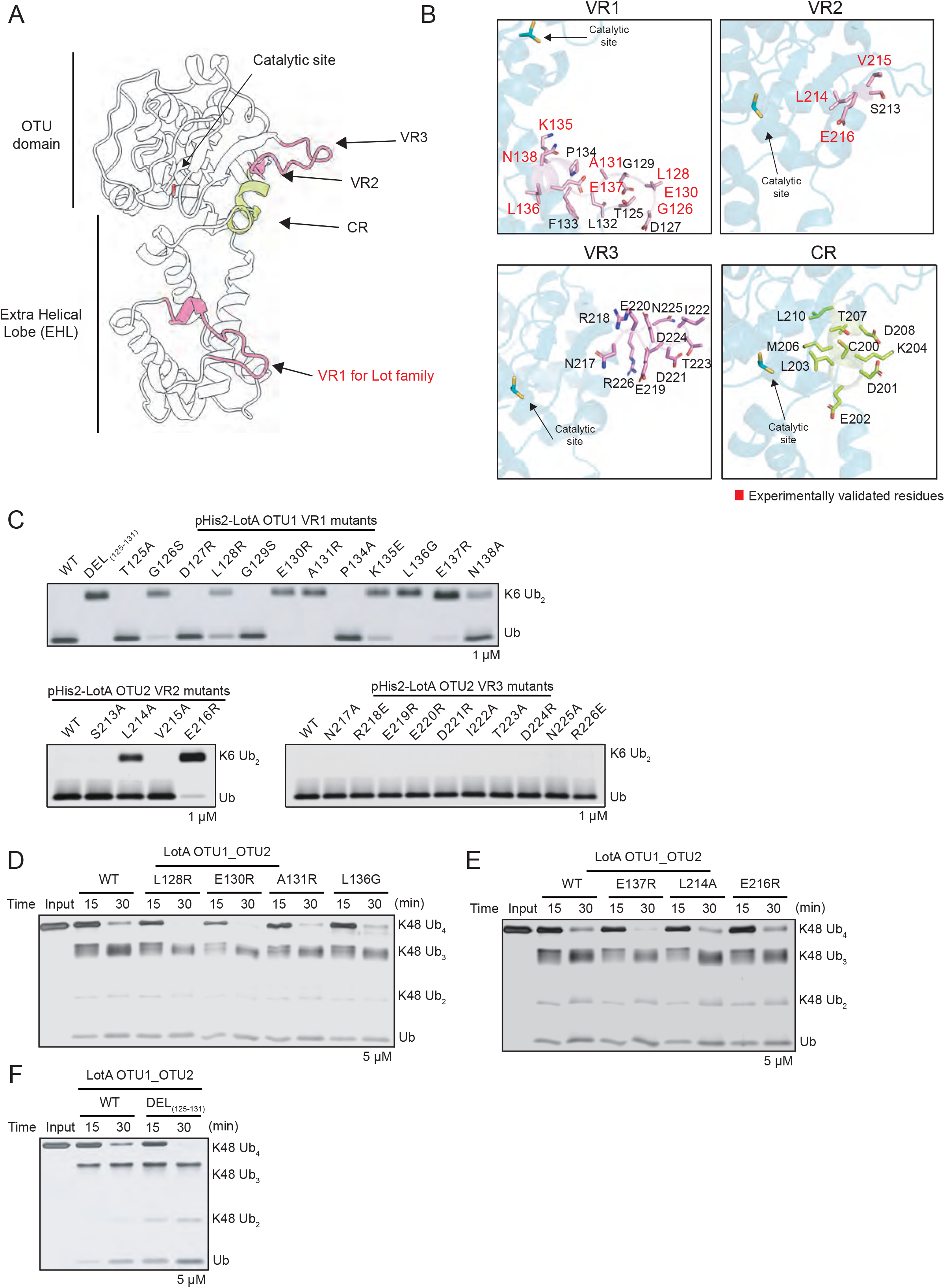
S1 ubiquitin-binding site of LotA OTU1. A. Cartoon representation of LotA OTU1. The constant region (CR, green) and three variable regions (VRs, pink) are compos part of the S1 ubiquitin-binding site. The EHL domain cotains the VR1 region of LOT DUB family. B. Close-up views of the VRs and CR of LotA OTU1. Three VRs (pink) and CR (green) residues are represented as stick model, and experimentally identified residues are highlighted as red color. C. K6-linked di-ubiquitin cleavage assay by LotA OTU1 wild-type and its mutants or deletion of VRs (VR1, VR2, and VR3) in LotA OTU1. Reactions were quenched at the indicated time-points and resolved by SDS-PAGE with silver staining. D-E. Poly-ubiquitin cleavage assay by LotA OTU1_OTU2 wild-type and VR1-mutants of LotA OTU1 as labeled. F. Gel-based K48-linked poly-ubiquitin cleavage assay by LotA OTU1_OTU2 wild-type and VR1eletion of LotA deletion (residues 125-131) mutant of LotA OTU1_OTU2.

Next, we investigated the effect of ubiquitin-binding sites of the OTU1 domain in the cooperative cleavage of K48-linked tetraubiquitin chains by OTU2 (Fig 7D and E). We introduced mutations in residues that we identified as ubiquitin-interacting residues on VR1 of the OTU1 domain into the LotA OTU1_OTU2 construct (LotA_7–544_). Surprisingly, some of these mutants (L128, E130, A131, E137) exhibited apparently different processing speeds for K48-linked tetraubiquitin cleavages, supporting the cooperation between the OTU1 and OTU2 domains. For further characterization, we deleted the VR1 loop of OTU1 from the OTU1_OTU2 construct and examined its catalytic activity (Fig 7F). The VR1 deletion mutant exhibited faster cleavage of the K48-linked chains than the WT, indicating the inhibitory roles of VR1 of OTU1 in K48-linked chain cleavage by OTU2. Overall, our results strongly support that the OTU1 and OTU2 domains of LotA simultaneously interact with each other to regulate the cleavage of K48-linked polyubiquitin chains.

## DISCUSSION

In this study, we investigated the structural and molecular basis of Lot DUBs. Structural analysis, including elucidation of the LotA OTU1 crystal structure, revealed that the EHL is found only in the Lot family (except LotA OTU1), shares the same structural topology, and provide the VR1, which acts as a ubiquitin-binding region. We also elucidated that the catalytic triad of the Lot family (except LotA OTU1) has a specific arrangement similar to that of A20-type OTU-DUBs. These unique molecular and structural characteristics categorize Lot DUBs as a unique OTU subfamily. Moreover, the activity of both LotA OTU1 and OTU2 domains is specific for certain ubiquitin chains, and LotA OTU1 assists LotA OTU2 in catalyzing the cleavage of K48- or K63-polyubiquitin chains. Thus, the structural and biochemical analyses presented in this study provide the conserved molecular basis of the Lot DUB family and unravel another aspect of the regulatory mechanism of DUBs.

DUBs, with linkage specificity, cleave specific polyubiquitin linkages. The ability to distinguish between different polyubiquitin linkages depends on various molecular determinants (Abdul Rehman *et al*., 2016; Faesen *et al*, 2011; Gersch *et al*, 2017; Kwasna *et al*., 2018; Mevissen *et al*., 2013). Our study revealed an additional noncanonical regulatory mechanism in DUBs. Unlike other DUBs that have additional ubiquitin-binding sites near the catalytic center or within the catalytic domain, LotA exhibits intramolecular cooperativeness between its two OTU domains. The OTU1 domain specifically cleaves K6-linked polyubiquitin chains, but it is also critical for the cleavage of K48-linked polyubiquitin chains by LotA OTU2. Moreover, structural restriction induced by the deletion of the linker between OTU1 and OTU2 resulted in reduced activity of LotA OTU2, indicating the cooperativeness between the two OTU domains of LotA.

Cooperative processing of K48-linked polyubiquitin chains by LotA is dependent on the length of the ubiquitin chain (Fig 2). We observed that intact LotA OTU1_OTU2 could not process K48-linked diubiquitin chains, but it could cleave K48-linked tetraubiquitin chains, indicating a length-dependent regulatory mechanism. Interestingly, recent studies have shown preferential cleavage of long K48-linked ubiquitin chains by the human DUBs MINDY1 and MINDY2 (Abdul Rehman *et al*, 2021; Abdul Rehman *et al*., 2016). MINDY1 has five different ubiquitin-binding sites (S1, S1′–S4′) that act as the determinants of the length of the K48-linked polyubiquitin chain. Similar to that of MINDY1, we observed that the S1 ubiquitin-binding site of the OTU1 domain affects the activity of the OTU2 domain. We observed an increased cleavage of K48-linked tetraubiquitin chains upon introduction of mutations in the VR1 of the OTU1 domain, which was identified as a K6-linked ubiquitin-binding site, suggesting an additional role of VR1 in the regulation of the cleavage of K48-linked polyubiquitin chains. Unlike MINDY1/2, which has five ubiquitin-binding sites, LotA has two catalytic domains and the ubiquitin-binding site on one catalytic domain regulates the other OTU domain. To further understand the intramolecular cooperation of LotA, structural analyses of K48-linked tetraubiquitin or longer chains with LotA OTU1_OTU2 are required. Nevertheless, the results of the present study add another aspect to the regulatory mechanisms of DUBs.

Accumulating evidence supports that *L. pneumophila* harbors a series of genes that alter the host ubiquitination system. Among them, many DUBs that cleave specific polyubiquitin chains have been identified. *Legionella* has an arsenal of DUBs that interrupt all possible polyubiquitin linkages (LotA (K6, K48), LotB (K63), LotC (K6, K11, K27, K29, K33, K48, K63), LotD (K6, K11, K48, K63), RavD (M1)) (Kubori *et al*., 2018; Liu *et al*., 2020; Ma *et al*., 2020; Schubert *et al*., 2020; Shin *et al*., 2020a; Takekawa *et al*., 2022; Wan *et al*., 2019). While previous studies have revealed the cellular localization or interacting host proteins of these DUBs, little is known about their physiological or pathological roles in infection. Therefore, systematic approaches are required to understand the underlying mechanism of DUBs in altering the host ubiquitination system.

## MATERIALS AND METHODS

### Protein expression and purification

All proteins used in this study were expressed and purified as previously reported (Bhogaraju *et al*., 2016; Qiu *et al*., 2016). Lpg2248 (LotA), Lpg1621 (LotB), Lpg2529 (LotC) were cloned into either pParallelHis2 or pParallelGST2 vector (Sheffield et al, 1999). BL21(DE3) Escherichia coli competent cells (NEB) were transformed with Lpg2248 (LotA) plasmids and T7 express Escherichia coli competent cells (NEB) were transformed with Lpg1621 (LotB) and Lpg2529 (LotC) plasmids, and both were grown in LB medium to an optical density (600 nm) of 0.6-0.8 at 37°C. After reaching an optical density of 0.6-0.8, protein expression was induced by the addition of 0.5 mM IPTG (isopropyl D-thiogalactopyranoside) for an additional 16 hr at 18°C and harvested. The cell pellet was resuspended in 50 mM Tris-HCl [pH 7.6], 150 mM NaCl, and 2 mM DTT (Lysis buffer) and lysed by sonication and centrifuged at 13,000 rpm to defecate the supernatant. The supernatant of GST-tagged protein was incubated for 2 hr with glutathione Sepharose 4B (Cytiva), and pre-equilibrated with 50 mM Tris-HCl [pH 7.6], 500 mM NaCl, and 2 mM DTT (Wash buffer), non-specific proteins were cleared by washing steps. Proteins were eluted with 50 mM Tris-HCl [pH 8.0], 50 mM NaCl, 2 mM DTT, and 15 mM L-Glutathione reduced (Elution buffer) and the buffer was exchanged to 50 mM Tris-HCl [pH 7.6] 150 mM NaCl, and 1 mM DTT (Storage buffer). For His-tagged proteins, the supernatant was incubated for 2 hr with TALON metal affinity resin (Takara) pre-equilibrated with 50 mM Tris-HCl [pH 7.6], 500 mM NaCl, 10 mM Imidazole, and 1 mM TCEP (Wash buffer). His-tagged proteins were eluted with 50 mM Tris-HCl [pH 7.6], 500 mM NaCl, 300 mM Imidazole, 1 mM TCEP (Elution buffer) and the buffer was exchanged to the storage buffer. For LotA_7-290_, glutathione beads were incubated with sfGFP-TEV protease (Wu et al, 2009) for 4 hr at 25°C and then overnight at 4°C, instead of using the elution buffer. GST-tagged cleaved proteins were exchanged buffer to 20 mM Tris-HCl [pH 8.0] 20 mM NaCl, and 1 mM DTT (IEX buffer A) and purified by anion-exchange chromatography on HitrapQ (Cytiva) with gradient elution using 20 mM Tris-HCl [pH 8.0], 1 M NaCl, and 1 mM DTT (IEX buffer B). Selected proteins were loaded onto size-exclusion column (Superdex 75 16/60, Cytiva), and pre-equilibrated with 50 mM Tris-HCl [pH 7.6], 50 mM NaCl, and 1 mM TCEP (SEC buffer). For crystallization, proteins were concentrated at 267 µM and stored.

### Crystallization

Purified and concentrated LotA_7-290_ were screened with the sitting drop vapor diffusion method, screened in a 96 well plate with 100 nl of protein and 200 nl of precipitant solution at 290 K. Initial crystals were found from solution containing 32% (w/v) PEG 4000, 0.1 M Tris-HCl [pH8.5], 0.8 M Lithium Chloride with 267 µM protein concentration. Diffraction-quality crystals were obtained from the optimized solution containing 35% (w/v) PEG 4000, 0.1 M Tris-HCl [pH7.8], 0.8 M Lithium Chloride with 267 µM protein concentration.

### Data collection, and structure determination

Crystals were cryo-protected using mother liquor solution containing 25% (v/v) ethylene glycerol. Diffraction data were collected at the Pohang Light Source (PLS) Beamline 11C. Initial data sets were processed using molecular replacement (McCoy et al, 2007) with Phaser in CCP4, using the prediction structure of AlphaFold2 Colab (Mirdita et al, 2022). Structure refinement and manual model building were performed with Coot and Phenix. Refine (Afonine et al, 2012; Emsley et al, 2010). All figures in this study were generated using PyMOL.

### Di-Ub panel cleavage assay

All di-ubiquitin chains used in this study were synthesized as previously described (El Oualid *et al*, 2010). To activate His-tagged LotA_7-290_, and LotA_7-290_ mutants, 3 µl of 1 µM DUBs were mixed with 12 µl of 25 mM Tris-HCl [pH 7.5], 150 mM NaCl, and 10 mM DTT (Activation buffer) and incubated 15 mins at 25°C. For His-tagged LotA_7-544_, LotA_294-544_, LotB, and LotC, 3 µl of 5 µM DUBs were mixed with 12 µl of activation buffer. For di-ubiquitin samples, 3 µl of di-ubiquitin chains (0.2 mg/ml) were mixed with 3 µl of 500 mM Tris-HCl [pH 7.5], 500 mM NaCl, and 50 mM DTT (Reaction buffer) and 12 µl of ultra-pure water. Reactions were initiated by mixed activated DUBs with di-ubiquitin, incubated at 37°C, and then samples were collected at the indicated time points. The reactions were quenched by the addition of 2X SDS-sample buffer, then analyzed by SDS-PAGE, and stained with a silver staining kit (Power Silver Stain Kit, ELPIS-Biotech).

### Poly-Ub panel cleavage assay

K48, K63-linked poly-ubiquitin chains used in this study were purchased from South Bay Bio. To activate Gst or His-tagged LotA_7-290_, LotA_7-544_, LotA_294-544_, 3 µl of 5 µM DUBs were mixed with 12 µl of 25 mM Tris-HCl [pH 7.5], 150 mM NaCl, and 10 mM DTT (Activation buffer) and incubated 15 mins at 25°C. For poly-ubiquitin samples, 3 µl of di-ubiquitin chains (0.2 mg/ml) were mixed with 3 µl of 500 mM Tris-HCl [pH 7.5], 500 mM NaCl, and 50 mM DTT (Reaction buffer) and 12 µl of ultra-pure water. Reactions were initiated by mixed activated DUBs with poly-ubiquitin, incubated at 37°C, and then samples were collected at the indicated time points. The reactions were quenched by the addition of 2X SDS-sample buffer, then analyzed by SDS-PAGE, and stained with a silver staining kit (Power Silver Stain Kit, ELPIS-Biotech).

### Ub-Prg, Ub-VME reactivity assay

K48, K63-linked di-ubiquitin Prg, and K48, K63-linked di-ubiquitin PA, and VME used in this study were synthesized as previously described (Flierman *et al*., 2016; Mulder *et al*., 2014). To activate His-tagged LotA_7-290_, LotA_294-544_, LotA_7-544_, 3 µl of 5 µM DUBs were mixed with 12 µl of 25 mM Tris-HCl [pH 7.5], 150 mM NaCl, and 10 mM DTT (Activation buffer) and incubated 15 mins at 25°C. For di-ubiquitin-Prg, VME probes, 3 µl of di-ubiquitin probe (0.5 mg/ml) were mixed with 3 µl of 500 mM Tris-HCl [pH 7.5], 500 mM NaCl, and 50 mM DTT (Reaction buffer) and 12 µl of ultra-pure water. Reactions were initiated by mixed activated DUBs with di-ubiquitin probe, incubated at 37°C, and then samples were collected at the indicated time points. The reactions were quenched by the addition of 2X SDS-sample buffer, then analyzed by SDS-PAGE, and stained with a silver staining kit (Power Silver Stain Kit, ELPIS-Biotech).

### Negative stained electron microscopy

LotA_7-544_ was diluted to 0.03 mg/mL in SEC buffer for negative staining. Four microliters of protein was loaded onto a glow-discharged CF200-CU carbon grid (Electron Microscopy Sciences, U.K.). After 5 min of incubation, grid-bound LotA7-544 was quickly washed with distilled water. To stain Lot7-544 molecules, 1% uranyl acetate was loaded onto the grid and incubated for 2 min. The stained grid was imaged using a JEOL-2100 plus 200 kV microscope (JEOL, Japan) equipped with a Rio9 detector (Gatan, U.S.A). Sixty micrographs were collected and processed using *CryoSAPRC* (Punjani et al, 2017). The extracted initial 7535 particles were used for 2-dimensional (2D) classification. The selected 1,594 particles of the 2D class are described as Fig 3A.

### Specimen preparation and cryo-EM data collection

The cryo-EM grid sample was prepared by diluting LotA_7-544_ from 2 mg/mL to 0.6 mg/mL in SEC buffer for cryo-EM. Quantifoil Au 1.2/1.3 200 mesh grids (Quantifoil, Germany) were glow discharged for 60 s using a Pelco easiGlow^TM^ Glow-Discharge Cleaning System (Ted Pella Inc., U.S.A.). Four microliters of LotA_7-544_ was loaded onto a glow-discharged grid and blotted for 3 s with blot force 3 using Vitrobot Mark IV (Thermo Fisher, U.S.A.).

C-clipped LotA_7–544_ cryo-grid samples were imaged using a Glacios transmission electron microscope (Thermo Fisher, U.S.A) equipped with a field-emission gun with a voltage of 200 kV and a Falcon IV detector (Thermo Fisher, U.S.A). LotA_7–544_ micrographs were obtained using a Falcon IV detector and automated data collection system EPU software (Thermo Fisher, U.S.A.). LotA_7–544_ was observed at a magnification of 150,000,× with a pixel size of 0.68 Å. Each micrograph was recorded for 5.68 s, and the total dose was 60.14 e^-^/Å^2^ (∼1 e^-^ /Å^2^ per frame); the dose rate was 10.59 e^-^/Å^2^/s. Lot_7–544_ was imaged with a defocus range from -1.0 μm to -1.8 μm and total fractionation as 60 frames per micrograph.

### Image processing

Computing resources were utilized in CMCI at SNU and GSDC at KISTI. For LotA7–544, 2,538 micrographs were obtained using the EPU software. Initial dataset processing was performed using the *CryoSPARC* software (Punjani *et al*., 2017). This dataset was motion-corrected to 6–55 movie frames per micrograph. Motion-corrected micrographs were used to estimate the contrast transfer function (CTF) using *CTFFIND4* (Rohou & Grigorieff, 2015). The CTF-estimated micrographs were filtered into the final 1,796 micrographs for data processing. We used 400 micrographs for full-set particle picking. Next, we performed deep-learning-based particle-picking software (Topaz train) (Bepler *et al*, 2020) using reference particles. A total of 497,860 particles were used for the two-round 2D classification. After 2D classification, the selected 176,036 particles were transferred from *CryoSPARC* to *RELION 3.1.3* (Scheres, 2012; Zivanov *et al*, 2018) and further processed. 2D classification was used for the selected particles. The final particles (113,381 particles) were used an initial 3-dimensional (3D) model and classification; 3D classification was classified into three classes and the model was refined. This model was deposited in the EMDB database (EMDB: 34350).

## DATA AVAIALBILITY

Structural information including atomic coordinates and structure factors for the LotA OTU1 (LotA_7-290_) structure is deposited in the Protein DataBank (http://www.rcsb.org/) under accession numbers 8GOK.

## ACKNOWLEDGMENTS

We thank Sagar Bhogaraju and Sissy Kalayil for the critical reading of the manuscript. We also thank Prof. Hyun Soo Cho for sharing synchrotron time. The authors also thank the staff at PAL 5C and 11C for their support during the crystallographic X-ray diffraction test and data collection. This work was supported by a National Research Foundation of Korea (NRF) grant funded by the Korean government (MSIT) (No. 2021R1C1C100396112, 2018R1A6A1A0302560722 and 2021M3A9I4021220) and the Yonsei University Research Fund of 2021 (2021-22-0050). GJvdHvN acknowledges funding by ZonMw (Off-road grant 451001026) and NWO (VIDI grant VI.VIDI.192.011).

## FIGURE LEGENDS

**Figure EV1. Deubiquitinase activities of LotA constructs**

A. K48-linked di-ubiquitin-Propargyl (Prg) activity-based probes (ABPs) were incubated as indicated time-points with LotA OTU constructs (OTU1, OTU2, OTU1_OTU2) and analyzed at indicated time-points on SDS-PAGE with silver staining.

B. K48-linked di-ubiquitin-Vinyl methyl ester (VME) activity-based probes (ABPs) with LotA OTU constructs (OTU1, OTU2, OTU1_OTU2) and analyzed at indicated time-points on SDS-PAGE with silver staining.

C. K63-linked di-ubiquitin-Propargyl (Prg) activity-based probes (ABPs) were incubated as indicated timepoints with LotA OTU constructs (OTU1, OTU2, OTU1_OTU2) and analyzed at indicated time-points on SDS-PAGE with silver staining.

D. K63-linked di-ubiquitin-Vinyl methyl ester (VME) activity-based probes (ABPs) were incubated as indicated timepoints with LotA OTU constructs (OTU1, OTU2, OTU1_OTU2) and analyzed at indicated time-points on SDS-PAGE with silver staining.

**Figure EV2. Negative-staining and Cryo-EM data processing workflow**

Negative-staining (A) and Single-particle Cryo-EM (B) data processing flow chart of LotA OTU1_OTU2.

**Figure EV3. Structural comparison of Legionella OTU deubiquitinases**

A. Secondary structure alignment of LotA OTU1 based on the crystal structure. The OTU domain of LotA is colored green, and EHL is colored blue.

B. Alphafold prediction structure of LotD. Catalytic cysteine and histidine of LotD are shown as ball and stick model. Additional EHL compared to hOTUD1 and vOTU is colored dark green.

C. Structure of EHL domain of LotD. The EHL domain of LOT DUB family shared roof-like folding.

**Figure EV4. Deubiquitinase activities of LotD**

A. Di-ubiquitin panel cleavage assay by wild-type LotD and catalytic triad residues (C13S, H256A, D6A) mutation LotD against eight di-ubiquitin linkages. Reactions were quenched at the indicated time-points and analyzed on SDS-PAGE with silver staining.

**Figure EV5. Structure determination process of LotA OTU1**

A. Structure determination process of LotA_7-290_ using molecular replacement with the Alphafold predicted structure as a template model.

B. Structural comparison of LotC apo (6YK8, pink) and LotC bound with ubiquitin (7BU0, blue). Structures were aligned by core OTU domain.

**Figure EV6. S1 ubiquitin-binding site of LOT DUB family**

A. Structural overlay of LotA OTU1 (cyan), LotA OTU2 (7F9X, orange), LotB (6KS5, gray), and LotC (7BU0, pink). Structures were aligned by their core OTU domains. Three VRs are highlighted as pink, and CR is colored in green.

B. The EHL domain of LotA OTU1 (blue), LotA OTU2 (7F9X, orange), LotB (6KS5, gray), and LotC (7BU0, pink). VR1 for LOT DUB family are concentrated in a helical loop region C-terminal to the EHL core helix.

C. Close-up views of the VRs and CR of LotB (6KS5). Three VRs (pink) and CR (green) residues are represented as stick model, and experimentally validated residues are highlighted as red color.

D. Close-up views of the VRs and CR of LotC (7BU0). Three VRs (pink) and CR (green) residues are represented as stick model, and experimentally validated residues are highlighted as red color.

E. Close-up views of the VRs and CR of LotA OTU2 (7F9X). Three VRs (pink) and CR (green) residues are represented as stick model, and experimentally validated residues are highlighted as red color.

